# Cross-modal Graph Contrastive Learning with Cellular Images

**DOI:** 10.1101/2022.06.05.494905

**Authors:** Shuangjia Zheng, Jiahua Rao, Jixian Zhang, Ethan Cohen, Chengtao Li, Yuedong Yang

**Affiliations:** Sun Yat-sen University, Galixir; Sun Yat-sen University; Galixir Technologies; IBENS, Ecole Normale Supérieure

**Author notes:** These authors contribute equally to this work (Shuangjia Zheng, Jiahua Rao).

## Abstract

Constructing discriminative representations of molecules lies at the core of a number of domains such as drug discovery, material science, and chemistry. State-of-the-art methods employ graph neural networks (GNNs) and self-supervised learning (SSL) to learn the structural representations from unlabeled data, which can then be fine-tuned for downstream tasks. Albeit powerful, these methods that are pre-trained solely on molecular structures cannot generalize well to the tasks involved in intricate biological processes. To cope with this challenge, we propose using high-content cell microscopy images to assist in learning molecular representation. The fundamental rationale of our method is to leverage the correspondence between molecular topological structures and the caused perturbations at the phenotypic level. By including cross-modal pre-training with different types of contrastive loss functions in a unified framework, our model can efficiently learn generic and informative representations from cellular images, which are complementary to molecular structures. Empirical experiments demonstrated that the model transfers non-trivially to a variety of downstream tasks and is often competitive with the existing SSL baselines, e.g., a 15.4% absolute Hit@10 gains in graph-image retrieval task and a 4.0% absolute AUC improvements in clinical outcome predictions. Further zero-shot case studies show the potential of the approach to be applied to real-world drug discovery.

## 1 Introduction

Learning discriminative representations of molecules is a fundamental task for numerous applications such as molecular property prediction, de novo drug design and material discovery [1]. Since molecular structures are essentially topological graphs with atoms and covalent bonds, graph representation learning can be naturally introduced to capture the representation of molecules [2, 3, 4, 5]. Despite the fruitful progress, graph neural networks (GNNs) are known to be data-hungry, i.e., requiring a large amount of labeled data for training. However, task-specific labeled data are far from sufficient, as wet-lab experiments are resource-intensive and time-consuming. As a result, training datasets in chemistry and drug discovery are typically limited in size, and neural networks tend to overfit them, leading to poor generalization capability of the learned representations.

Inspired by the fruitful achievements of the self-supervision learning in computer vision [6, 7] and natural language processing [8, 9], there have been some attempts to extend self-supervised schemes to molecular representation learning and lead to remarkable results [10, 11, 12, 13, 14, 15]. The core of self-supervised learning lies in establishing meaningful pre-training objectives to harness the power of large unlabeled data. The pre-trained neural networks can then be used to fine-tune for small-scale downstream tasks.

However, pre-training on molecular graph structures remains a stiff challenge. One of the limitations of current approaches is the lack of domain knowledge in chemistry or chemical synthesis. Recent studies have pointed out that pre-trained GNNs with random node/edge masking gives limited improvements and often lead to negative transfer on downstream tasks [10, 16], as the perturbations actions on graph structures can hurt the structural inductive bias of molecules. Furthermore, molecular modeling tasks often require predicting the binding/interaction between molecules and other biological entities (e.g., RNA, proteins, pathways), and further generalizing to the phenotypic/clinical outcomes caused by these specific bindings [17]. Self-supervised learning methods that solely manipulate molecular structures struggle to handle downstream tasks that involve complex biological processes, limiting their practicality in a wide range of drug discovery applications.

To this end, we propose using high-content cell microscopy images to assist in learning molecular representation, extending the molecular representation beyond chemical structures and thus, improving the generalization capability. High-content cell microscopy imaging (HCI) is an increasingly important biotechnology in recent years in the domain of drug discovery and system biology [18, 19]. As shown in Figure 1, small molecules enter into cells and affect their biological functions and pathways, leading to morphological changes in cell shape, number, structure, etc., that are visible in microscopy images after staining. Analysis and modeling based on these high-content images have shown great success in molecular bioactivity prediction [20], mechanism identification [21], polypharmacology prediction [22], etc. The stained cell images contain rich morphological information that reflects the biological changes induced by chemical structures on cell cultures. Thus, we hypothesize that this phenotypic modality has complementary strengths with molecular structures to make enhanced representations and thus benefit the downstream tasks involved in intricate biological processes.

**Figure 1:**
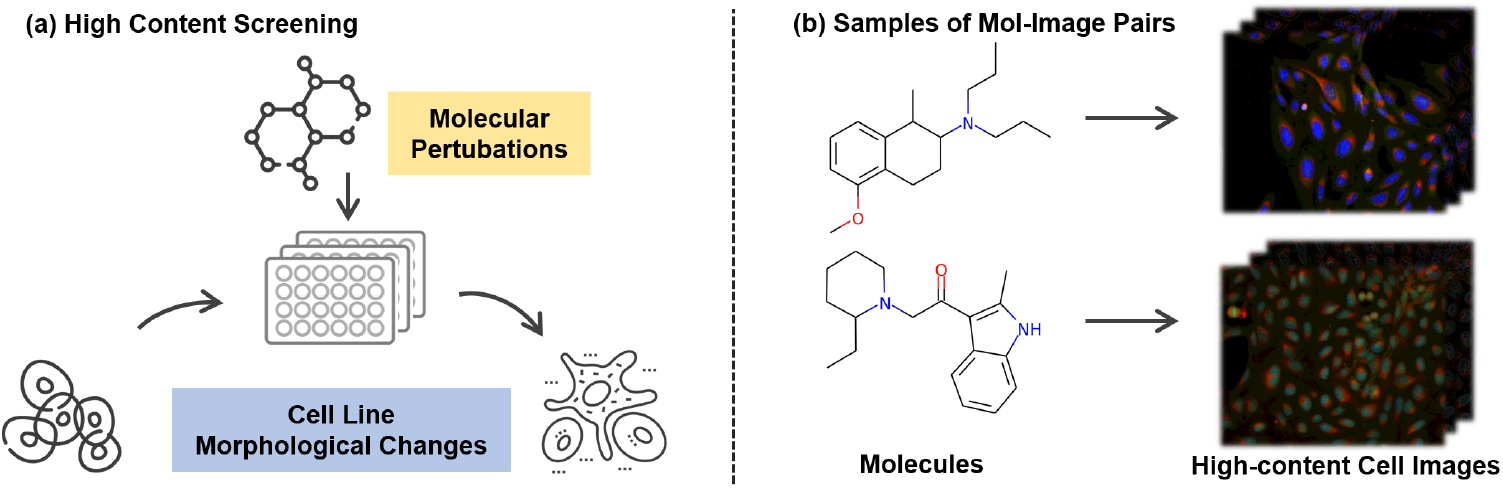
(a) Illustration of high-content screening experimental protocol. (b) Samples of molecule-images pairs. Note that one molecule produces multiple images.

However, building connections between molecular structures and high-content cellular images is a challenging task that highlights representative, unsolved problems in cross-modal learning. The first challenge comes from the unclear fine-grained correspondence between molecules and images. Unlike conventional cross-modal paired data such as caption and picture, the patterns of the molecule are not directly reflected in the cellular image, thus preventing us from using traditional cross-modal encoders for alignment. The second challenge arises from the noise and batch effect of the cellular data. For example, cellular images obtained from the same molecule can vary considerably. Existing cross-modal pre-training objectives may overfit the noisy images and decrease the model’s generalization ability.

Herein, we propose Molecular graph and hIgh content imaGe Alignment (MIGA), a novel cross-modal graph-and-image pre-training framework to address the above issues. We first encode the molecule and cell microscopy images independently with a molecular graph encoder and an image encoder. Then we align the graph embeddings with the image embeddings through three contrastive modules: graph-image contrastive (GIC) learning, masked graph modeling (MGM) and generative graph-image matching (GGIM). Specifically, (i) GIC encourages high similarity between the latent embeddings of matched graph-image pairs while pushing those of non-matched pairs apart; (ii) MGM, a local cross-modal module that utilizes both the observed (unmasked) graph and the cell image to predict the masked molecular patterns and (iii) GGIM aims to learn an effective cross-modal representation by improving graph-image matching accuracy and cross-modality generalization ability. The three modules are complementary and thus the combination of these modules can 1) make it easier for the encoders to perform cross-modal learning by capturing structural and localized information; (2) learn a common low-dimensional space to embed graphs and images that are biologically meaningful. Enabled by the massive publicly available high-content screening data [23], we establish a novel cross-modal benchmark dataset that contains 750k molecular graph-cellular image pairs. To evaluate models on this benchmark, we propose a new biological meaningful retrieval task specific to graph-image cross-modal learning. We also include existing clinical outcome prediction and property prediction task to further assess the learned representations. Extensive experiments demonstrate that the cross-modal representations learned by our proposed model, MIGA, can benefit a wide-range of downstream tasks that require extensive biological priors. For example, MIGA achieves a 15.4% absolute Hit@10 gain in graph-image retrieval task and a 4.0% absolute AUC improvement in clinical outcome predictions, respectively, over existing state-of-the-art methods.

## 2 Related work

### Self-supervised learning on graphs

Graph self-supervised pre-learning attempts to obtain supervision in unlabelled data to learn meaningful representations that can be further transferred to downstream tasks [12, 10, 24, 11, 13, 14, 15, 25, 26, 27, 16]. In general, these methods fall into two categories: contrastive-based methods and generative-based methods [28]. The former aims to generate different views from the original graph and perform intra-graph contrastive learning [10, 11, 12, 13, 15], while the latter ones are trained in a supervised manner to generate masked sub-patterns or attributes at the inter-graph level [10, 14, 26]. These approaches achieve remarkable performance on molecular graph representation tasks, but lack the ability to predict the complex properties involved in intricate biological processes. Different from the previous works, our method leverages pairwise data from molecular and cellular image to improve the biological perception of the learned molecular representation in a novel cross-modal pre-training framework.

### Cross-modal pre-training

Pre-training strategies for multi-modal tasks have attracted massive attention, with most of these efforts targeting Visual-Language representation learning. Most of them can be grouped into two categories. The first category is to use multi-modal encoders to capture the interaction between image and text embeddings [29, 30, 31, 32, 33, 34]. Approaches in this category achieve remarkable performance, but most of them require high-quality images and pre-trained object detectors. The other category focuses on learning independent decoders for different modalities [35, 36, 37]. For instance, CLIP [35] learns pairwise relationships between language and images by performing pre-training on a large amount of web data using independent encoders connected by contrastive losses. Our method shares similar spirits with the second category, but with clear differences. Unlike the comprehensible correspondence between images and text, our modalities are graph structure and cell image. Their connection is implicit and lacks a fine-grained one-to-one correspondence. This also led us to focus more on the overall similarity and the noise reduction.

## 3 Methods

At the core of our method is the idea of infusing structural representations with biological perception by building the connections between molecular graph and their induced morphological features.

To achieve this, as illustrated in Figure 2, given pairs of graph and image, we employ two independent encoders (a GNN *f*^*g*^ and a CNN *f*^*i*^) to produce the representations of a molecular graph *G* and a cellular image *I*, and align them with inter- and intra-contrastive losses. This cross-modal framework pulls the matched graph-image pairs together and contrasts the unmatched pairs apart. After pretraining, we use the output representations to make zero-shot predictions or fine-tune the networks on downstream tasks. This cross-modal learning process can also be interpreted as morphological information being passed through the convolutional neural network to the graph neural network in a knowledge distillation manner [38].

**Figure 2:**
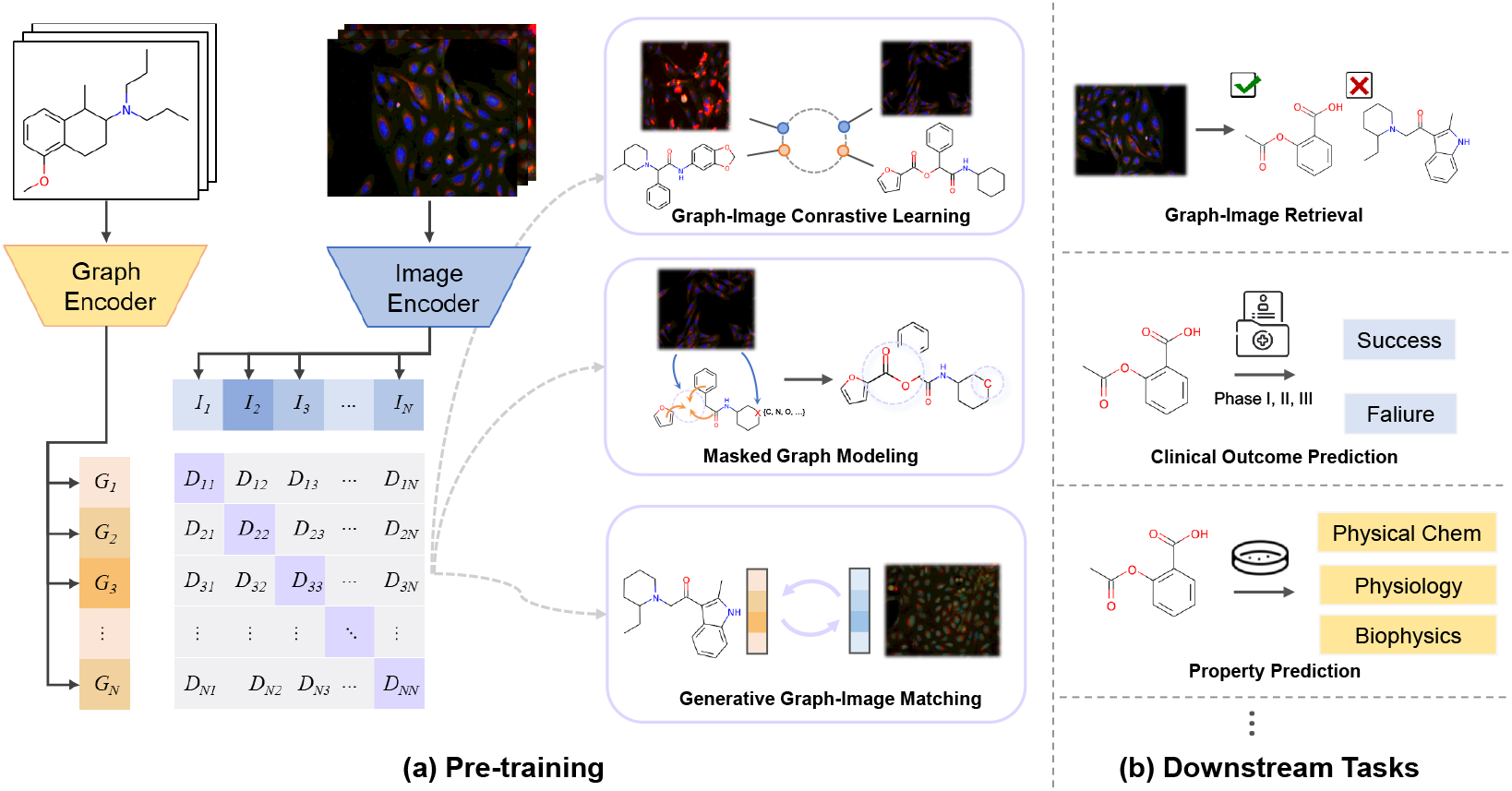
Overview of our method, MIGA. Molecular graph and cellular image representations are jointly learned from pairwise data. We used three complementary contrastive objectives to perform cross-modal pre-training. The learned representations can be used for graph-image or graph-only downstream task transfer.

In the following sections, we first introduce the structural and image encoders for cross-modal learning (section 3.1). Then we depict the details of contrastive training objectives (section 3.2), followed by the experimental setting (section 4.1) and results (section 4.2).

### 3.1 Structure and Image Encoders

#### Structural encoder

A compound structure can be represented as an attributed graph *G* = (𝒱, ℰ), where |𝒱| = *n* denotes a set of *n* atoms (nodes) and |ℰ| = *m* denotes a set of *m* bonds (edges). We represent 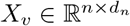 for the node attributes with *d*_*n*_ as the the feature dimension of node and 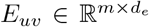 for the edge attributes with *d*_*e*_ as the feature dimension of edge. A graph neural network (GNN) *f* ^*g*^ learns to embed an attributed graph *G* into a feature vector *z*_*G*_. We adopt the Graph Isomorphism Network (GIN) from [3], where the node and edge attributes are propagated at each iteration. Formally, the k-th iteration of a GNN is:

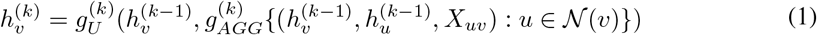

where 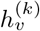 are the representation of node *v* at the *k*-th layer, 𝒩(*v*) is the neighbourhood set of node 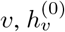 is initialised with *X*_*v*_ encoding its atom properties. 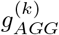 stands for the aggregation function and 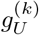 stands for the update function. After *K* graph convolutions, *h*^*K*^ have captured their *K*-hop neighbourhood information. Finally, a readout function is used to aggregate all node representations output by the *K*-th GNN layer to obtain the entire molecule’s representation *z*_*G*_:

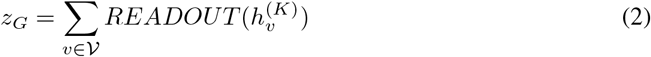

#### Image encoder

For the cellular image *I*, we first use residual convolutional neural networks (ResNet-34) [39] as the basic feature extractor *f*^*i*^ because it is lightweight, widely-adopted and have proven performance. We also implement DenseNet [40], EfficientNet [41] and the recently proposed Vision Transformer (ViT) [42] as comparisons. We strictly follow their original implementation with a small modification, namely adding an additional projection layer before the final output. These models are initialized with weights pre-trained on ImageNet [43]. Each input image *I* is encoded into a one-dimensional feature vector *z*_*I*_ for further fusion.

### 3.2 Pre-training framework

We pre-trained MIGA through three contrastive objectives: graph-image contrastive (GIC) learning, masked graph modeling (MGM) and generative graph-image matching (GGIM).

#### Graph-image contrastive (GIC) learning

aims to pull embeddings of the matched molecule-image pairs together while pushing those of unmatched pairs apart by maximizing a lower bound on the mutual information (MI) between the molecular graph and cellular image for the positive pairs. We achieve this by minimizing a symmetric InfoNCE loss [44] to maximize a lower bound on MI(*G*; *I*). Formally, the graph-image contrastive loss is defined as:

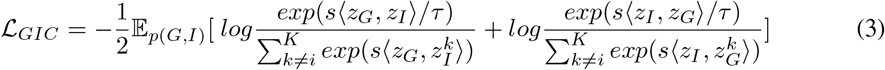

Here, the similarity function *s* ⟨*z*_*G*_, *z*_*I*_⟩ = *f*_*v*_(*z*_*G*_) · *f*_*w*_(*z*_*I*_)*/*(∥*f*_*v*_(*z*_*G*_)∥∥*f*_*w*_(*z*_*I*_)∥), where *f*_*v*_ and *f*_*w*_ are two linear projection layers that embed representations to a common space. *τ* is a temperature parameter, *K* is a set of negative image samples that not matched to *G*.

#### Masked graph modeling (MGM)

To simultaneously leverage the intra-molecular graph information and strengthen the interaction between molecule and image, we further utilize both the image and the partial molecular graph to predict the masked sub-patterns. Following the masking strategies of [10], we randomly mask the atom/bond attributes and constructed the context graphs for each molecular graph. We use the surrounding partial graph structures along with the corresponding image information to predict the masked attributed subgraphs and the corresponding attributes. Our goal is to pre-train molecular GNN *f*^*g*^ that can not only learn the context information of atoms in similar structures, but also capture domain knowledge by learning the regularities of the node/edge attributes distributed over graph structure. Herein, we defined the masked molecular graph as *G*^*msk*^ and the observed (unmasked) graph as *G*^*obs*^. Therefore, the training graph-image pair (*G, I*) have been transformed to (*G*^*msk*^, *G*^*obs*^, *I*) and the objective of MGM is:

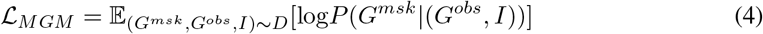

where (*G*^*msk*^, *G*^*obs*^, *I*) is a randomly sampled graph-image pair from the training set *D*. Thus, the MGM training is equivalent to optimize the following equation:

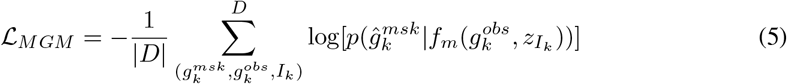

where the 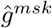 is the prediction from the observed graph *g*^*obs*^ and image embedding *z*_*I*_. The function *f*_*m*_ is the molecule attributes and context graphs prediction model. Our MLM training minimized by a cross-entropy loss because *g*^*msk*^ is a one-hot vocabulary distribution where the ground-truth attributes/subgraphs has a probability of 1.

#### Generative Graph-image matching (GGIM)

aims to learn an effective cross-modal representation by improving graph-image matching accuracy and cross-modality generation ability. Inspired by [45], we firstly utilize a cross-modal encoder *f*_*c*_ to fuse two separate unimodal representations and produce a joint representation of the graph-image pair, and append a multilayer perceptron followed by softmax to predict the matching probability of *p*^*m*^ = *f*_*c*_(*z*_*G*_, *z*_*I*_). This can be formulated as:

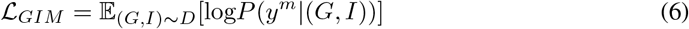

where *y*^*m*^ is the ground-truth label representing whether the graph and the image is matched or not matched. The expected log-likelihood function could be defined as:

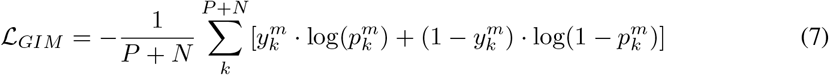

where *P* denotes the number of positive graph-image pairs and *N* denotes the number of negative pairs.

Different from most cross-modal pre-training works that have obvious correspondence features (e.g. a word in the text is related to an entity in the image), the correspondences of the graph-image pairs being used for pre-training are implicit. Moreover, there may exist other images different from the ground-truth that reflects the molecular perturbation equally well (or better) because of the batch effect [46]. Motivated by recent success in generative contrastive learning [47, 27], we further employ variational auto-encoders (VAE) [48] as generative agents to reduce noise from experiments and enhance the ability of cross-modal generalization. In particular, the generative agents are asked to recover the representation of one modality given the parallel representation from the other modality. Herein, we performed cross-modal generation from two directions including generating cellular images from matching topological graphs and generating graphs from images. This generative graph-Image matching (GGIM) loss function can be formulated as:

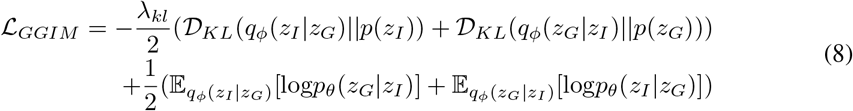

where *λ*_*kl*_ is the hyperparameter balancing the reconstruction loss. *p*(*z*_*I*_) and *p*(*z*_*G*_) is the prior of image embedding and graph embedding, respectively, and *q*_*ϕ*_(*z*_*G*_|*z*_*I*_), *q*_*ϕ*_(*z*_*I*_|*z*_*G*_) is the corresponding posterior. Finally, the full contrastive objective of MIGA is:

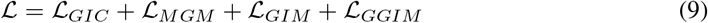

## 4 Experiments

### 4.1 Datasets and Tasks

#### Pre-training Dataset

We perform our experiments on the Cell Painting dataset CIL introduced by [19, 23]. The dataset originally consists of 919,265 cellular images collected from 30,616 molecular interventions. Each image contains five color channels that capture the morphology of five cellular compartments: nucleus (DNA), Endoplasmic reticulum (ER), nucleolus/cytoplasmic RNA (RNA), F-actin cytoskeleton (AGP) and Mitochondria (Mito). A molecular interventions is photographed from multiple views in an experimental well and the experiment was repeated several times, resulting in an average of 30 views for each molecule. In order to keep the data balanced, we restricted each molecule to a maximum of 30 images and removed the untreated reference images, resulting in a cross-modal graph-image benchmark containing 750K views. We refer to this benchmark as CIL-750K. Detailed pre-process procedure and analysis of pre-train dataset can be found in Appendix A.

#### Downstream Tasks

We evaluate the pre-trained model in three downstream tasks: graph-image retrieval, clinical outcome prediction and molecular property prediction.

- **Graph-Image Retrieval** contains two tasks: (1) graph as query and image as targets (Image retrieval); (2) image as query and graph as targets (graph retrieval). The goal of this task is to demonstrate whether the learned embeddings are able to preserve the inherent relations among corresponding graph-image pairs. We randomly split the CIL-750K dataset into a training set of 27.6K molecules corresponding to 680K images, and hold out the remaining of the data for testing. The held-out data consists of 3K molecules and the corresponding 50K images. We formulate the retrieval task as a ranking problem. In the inference phase, given a query molecular graph in the held-out set, we take images in the held-out set as a candidate pool and rank candidate images according to the L2 distance between the image embeddings and the molecular embeddings, and vice versa. The ranking of the ground-truth image/graph can then be used to calculate AUC, MRR (mean reciprocal rank), and Hit@1, 5, 10 (hit ratio with cut-off values of 1, 5, and 10).
- **Clinical Trial Outcome Prediction** aims to predict the clinical trial outcome (success or failure) of drug molecules. This task is extremely challenge as it requires external knowledge on system biology and trial risk from the clinical record. We use the Trial Outcome Prediction (TOP) benchmark constructed by [49] for model evaluation. After dropping the combination mediation, the benchmark contains 944, 2865 and 1752 molecules for Phase I, II, III tasks, respectively. We follow the data splitting proposed by [49] and employ Precision-recall area under the curve (PR-AUC) and Area under the receiver operating characteristic curve (ROC-AUC) to measure the performance of all methods.
- **Molecular Property Prediction** We further evaluate MIGA on six molecular property datasets: HIV, Tox21, BBBP,ToxCast, ESOL and Lipophilicity. These datasets are introduced by [1] and further benchmarked by OGB community[50] for low-resource graph representation learning. Each data set contains thousands of molecular graphs as well as binary/scalar labels indicating the property of interest. We follow the OGB [50] setting and adopt the scaffold splitting strategy with a ratio for train/valid/test as 8:1:1 during fine-tuning.

### 4.2 Results

#### 4.2.1 Graph-Image Retrieval

##### Baselines

Since we present improvements to pre-training models, we primarily compare our approach to the existing state-of-the-art graph SSL method, GraphCL [12] and cross-modal pre-learning methods, CLIP [35] and ALIGN [36]. As none of them were specifically designed for graph-image cross modal learning, we adapted their encoders but followed their loss functions and tricks to perform pre-training on CIL-750K. We then evaluate the effectiveness of the proposed pre-training objectives (i.e., GIC, MGM, GIM, GGIM) with four variants of our method. For each baseline, we use the pre-trained model to output embeddings of molecular graphs and celluar images, then rank the candidate pool based on their similarity. We also include random initialization and feature engineering-based representation (ECFP [51] + CellProfiler [52]) as baselines to demonstrate the challenge of the task. More details and hyperparameters have been attached in Appendix B.

##### Result on CIL-750K held-out set

Results of graph-image retrieval tasks are shown in Table 1. It is clear that our method MIGA significantly outperform baseline methods including ECFP+CellProfiler, GraphCL, CLIP and ALIGN. For example, MIGA achieves 12.9% absolute MRR gain and 15.4% absolute Hit@10 gain over the best baseline ALIGN on image retrieval task. These improvements are consistent in graph retrieval task, proving that the proposed method has superior performance in graph-image cross modal learning. Compared to the basic loss (GIC), adding MGM and GIM both substantially improves the pre-trained model’s performance across two tasks. The proposed GGIM further enhances the model by reducing the noisy and building implicit connection between two modality.

**Table 1:** Graph-image retrieval tasks on held-out set of CIL-750K. MIGA and its variants are compared with adapted pre-training methods, GraphCL [12], CLIP[35], ALIGN[36], along with random initialization and a feature-based method ECFP [51] + CellProfiler [52]. The average of MRR, AUC, Hit@1, Hit@5 and Hit@10 are reported. The best and second best results are marked **<u>bold</u>** and **bold**, respectively.

| Task Metrics | Image Retrieval |  |  |  |  | Graph Retrieval |  |  |  |  |
| --- | --- | --- | --- | --- | --- | --- | --- | --- | --- | --- |
|  | MRR | AUC | Hit@1 | Hit@5 | Hit@10 | MRR | AUC | Hit@1 | Hit@5 | Hit@10 |
| Random Init | 0.051 | 0.500 | 0.010 | 0.050 | 0.100 | 0.051 | 0.500 | 0.010 | 0.050 | 0.100 |
| ECFP+CellProfiler | 0.052 | 0.511 | 0.010 | 0.049 | 0.100 | 0.053 | 0.500 | 0.011 | 0.050 | 0.100 |
| GraphCL | 0.247 | 0.818 | 0.106 | 0.391 | 0.558 | 0.279 | 0.835 | 0.132 | 0.433 | 0.616 |
| CLIP | 0.265 | 0.890 | 0.105 | 0.437 | 0.635 | 0.327 | 0.878 | 0.171 | 0.515 | 0.672 |
| ALIGN | 0.288 | 0.821 | 0.148 | 0.434 | 0.594 | 0.379 | 0.810 | 0.214 | 0.581 | 0.713 |
| GIC | 0.288 | 0.847 | 0.148 | 0.434 | 0.594 | 0.280 | 0.907 | 0.125 | 0.447 | 0.652 |
| GIC+GIM | 0.303 | 0.876 | 0.145 | 0.485 | 0.660 | 0.337 | 0.927 | 0.176 | 0.523 | 0.693 |
| GIC+MGM | **0.409** | 0.913 | **0.244** | 0.612 | 0.741 | 0.405 | 0.931 | 0.244 | 0.598 | 0.737 |
| GIC+MGM+GIM | 0.401 | **0.926** | 0.230 | **0.616** | **0.743** | **0.433** | **0.935** | **0.275** | **0.628** | **0.739** |
| MIGA (FULL) | **0.417** | **0.936** | **0.248** | **0.623** | **0.748** | **0.413** | **0.940** | **0.249** | **0.614** | **0.739** |

##### Case study for image retrieval

We randomly select 20 pairs in the held-out set for case study. Figure 3 shows the results of four molecules, while the full version is attached in Appendix C. The result demonstrates that our model is more powerful in prioritizing the images that are similar to the ground-truth compared to ALIGN.

**Figure 3:**
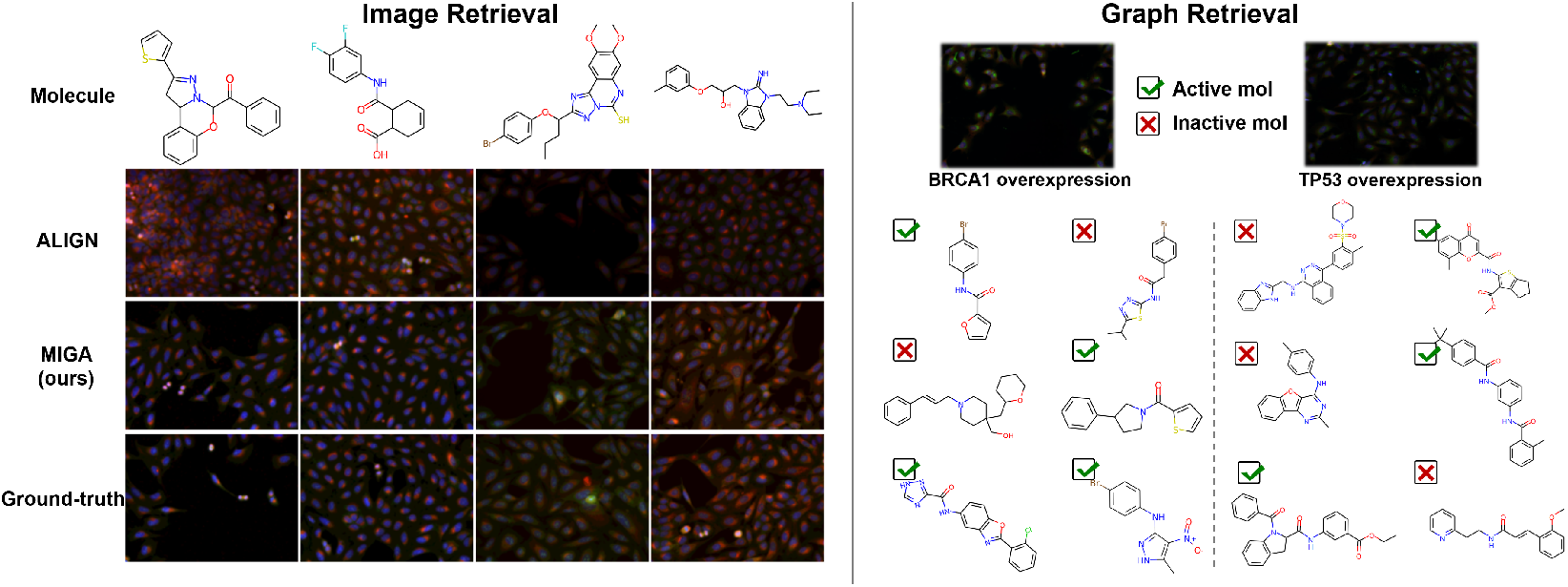
Case study on image retrieval task (left) and zero-shot image retrieval task (right). The images retrieved by our method and baseline method are shown on the left. Right side shows the cells induced by cDNA and our model can find diverse molecules that have similar functions to cDNA interventions (ticked). The full version is in Appendix D.

##### Case study for zero-shot graph retrieval

Of more practical value in the real world is the use of cellular images induced by non-small molecule interventions (e.g. cNDA [53]) to recall small molecules with similar effects. We collect 6 sets of cellular image induced by cDNA interventions for specific genes from [53]. We use ExCAPEDB [54] to retrieve active (agonists) and inactive molecules of these six genes as the candidate pools. Each gene has 20 active and 100 inactive molecules. Specifically, the mean Hit@10 for MIGA was 0.49 ± 0.05, much higher than the random enrichment 0.167 (20/120), demonstrating the ability of our method to generalize to new datasets in a zero-shot manner. This result also indicates the practical potential of our method in virtual screening and drug repurposing. Figure 3 shows the results of two cellular images with cDNA-induced gene over-expression, the top-6 molecules ranked by MIGA are shown, see Appendix D for the full version.

#### 4.2.2 Clinical trial outcome prediction

##### Baselines

Since clinical prediction is a single-modality task, we compare the proposed method with the state-of-the-art SSL methods, ContextPred [10], GraphLoG [11], GROVER [14], GraphCL [12] and JOAO [13]. We also included three machine learning-based methods (RF, LR, XGBoost) and a knowledge-aware GNN model HINT as listed in [49].

##### Results

Results are summarized in Table 2. Our best model MIGA improves PR-AUC by an average of 1.67% and AUC by an average of 3.97% on the three clinical outcome prediction datasets, respectively, compared with the best SSL baseline JOAO. This is consistent with our hypothesis that phenotypic features can help predict tasks that involve complex biological processes.

**Table 2:**
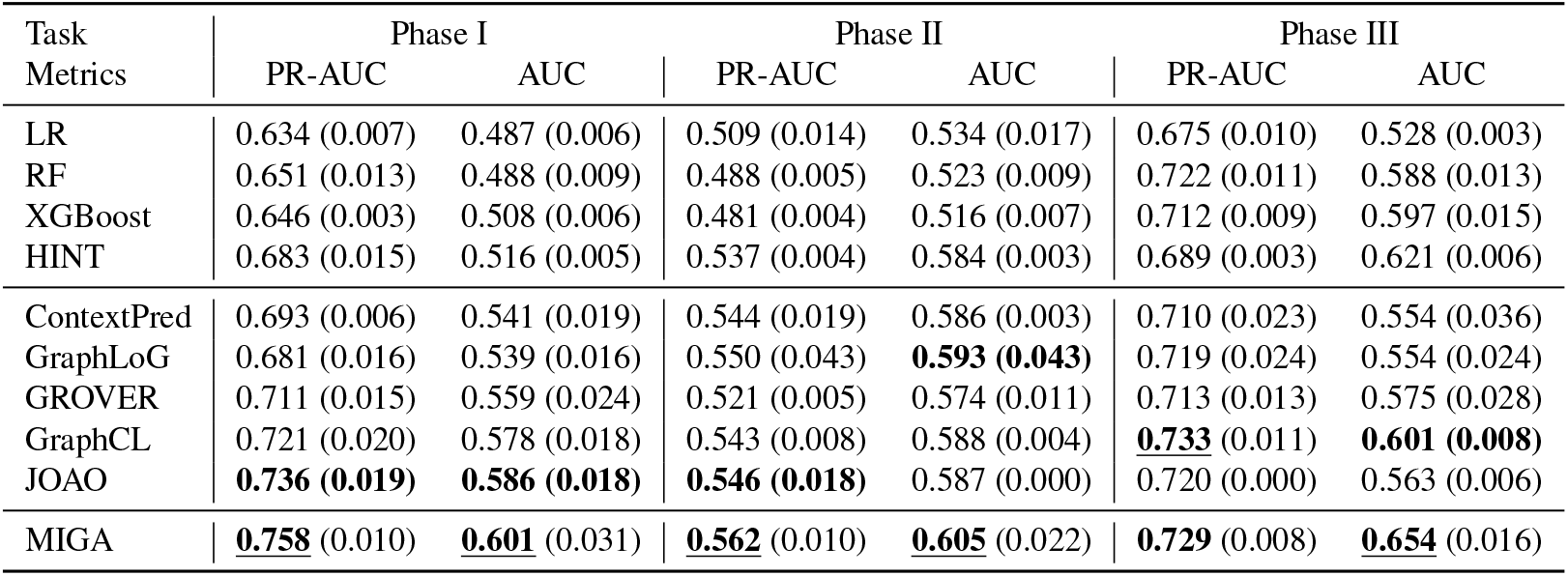
Performance comparison of MIGA and several baseline approaches for phase-level-outcome predictions on TOP dataset. We report the mean (and standard deviation) PR-AUC and ROC-AUC of five times for clinical trial outcome prediction. The best and second best results are marked **<u>bold</u>** and **bold**, respectively.

#### 4.2.3 Molecular property prediction

##### Baselines

We compare the proposed method with the SOTA SSL methods including EdgePred[55], ContextPred [10], AttrMasking[10], GraphCL [12], InfoGraph [25], GROVER[14], GraphLoG[11], GraphCL[12], JOAO[13]. All the SSL models are pre-trained using released source code on CIL-750K data set. For fine-tuning, we follow the same setting with [12, 13].

##### Results

Results are summarized in Table 3. As we can see, MIGA performs the best on 5 out of 6 datasets and outperforms the existing SSL methods in the average performance by a large margin. The slightly lower performance in ESOL is expected as this property related to molecule’s intrinsic characteristic, e.g., hydrophilicity and hydrophobicity. It does not rely on biological processes and phenotpyic changes. Even though, MIGA will not cause negative transfer compared to non-pretrain baseline.

**Table 3:** Comparison of SSL baselines against MIGA on six OGB datasets. Mean ROC-AUC and Root Mean Squared Error (RMSE) (with the SD) of 5 times independent test are reported. The best and second best results are marked **<u>bold</u>** and **bold**, respectively.

| Dataset | Classification (AUC) |  |  |  |  | Regression (RMSE) |  |  |
| --- | --- | --- | --- | --- | --- | --- | --- | --- |
|  | HIV | Tox21 | ToxCast | BBBP | Avg. | ESOL | Lipo | Avg. |
| Non-pretrain | 70.30 (0.51) | 68.90 (0.80) | 58.60 (1.20) | 65.40(2.4) | 65.80 | 1.278 (0.24) | 0.744 (0.14) | 1.011 |
| ContextPred | 74.17 (1.33) | 71.44 (0.11) | 60.05 (0.15) | 69.87 (0.99) | 68.88 | 1.141 (0.03) | 0.724 (0.02) | **0.933** |
| AttrMask | 75.55 (1.00) | 74.58 (0.66) | 59.51 (0.36) | 68.88 (2.65) | 69.63 | 1.194 (0.04) | 0.736 (0.02) | 0.965 |
| EdgePred | 72.53 (1.20) | 68.86 (0.38) | 57.39 (0.80) | 63.45 (1.39) | 65.56 | 1.146 (0.05) | 0.751 (0.02) | 0.949 |
| InfoGraph | **76.22 (0.24)** | 69.22 (0.78) | 59.87 (0.35) | 63.75 (1.52) | 67.27 | 1.242 (0.01) | 0.725 (0.01) | 0.984 |
| GraphLoG | 73.29 (2.64) | 69.80 (0.41) | 59.22 (1.05) | 68.43 (2.86) | 67.68 | 1.194 (0.02) | 0.766 (0.01) | 0.980 |
| GraphCL | 74.85 (1.71) | 74.19 (0.43) | 61.37 (0.10) | 66.13 (1.68) | 69.14 | 1.151 (0.04) | 0.745 (0.02) | 0.948 |
| GROVER | 74.35 (0.92) | 74.02 (0.79) | 61.30 (0.13) | **69.88 (0.58)** | **69.89** | 1.199 (0.02) | **0.721 (0.01)** | 0.960 |
| JOAO | 74.91 (0.66) | 74.60 (0.49) | **61.62 (0.37)** | 68.33 (0.58) | 69.87 | **1.117 (0.05)** | 0.753 (0.02) | 0.935 |
| MIGA | **76.38 (0.55)** | **75.23 (0.71)** | **62.34 (0.23)** | **71.52 (0.43)** | **71.37** | **1.123 (0.01)** | **0.717 (0.00)** | **0.919** |

### 4.3 Ablation study

Table 4 shows the effect of the number of view per molecule perform on graph retrieval task. We notice that with less than 10 views, the more views involve in pre-training the better the results would be, but using all views (on average 25 views per molecule) does not achieve more gains. We attribute this phenomenon to the batch effect of the cellular images, where images from different wells can vary considerably because of the experimental errors and thus mislead the model.

**Table 4:**
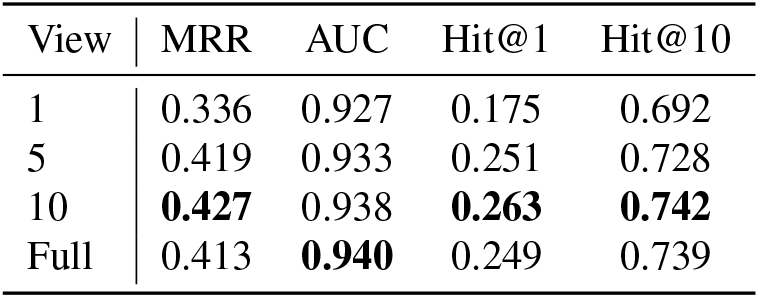
Effect of # view per molecule.

Table 5 studies the impact of CNN architecture choices on the graph retrieval task. Due to the relatively small amount of data, we use small CNN variants (ResNet34[39], EfficientNet-B1[41], DenseNet121[40] and ViT_tiny[42]) for evaluation. We note that small CNN models such as ResNet, EfficientNet and DenseNet achieve comparable performance while bigger models like ViT do not show superiority. We assume this is because these models are pre-trained on non-cellular images, so heavier models do not necessarily lead to gains on cellular tasks.

**Table 5:**
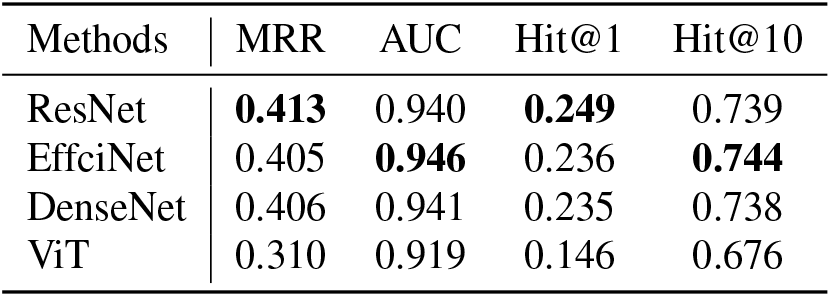
Effect of CNN architecture.

## 5 Conclusions

In this paper, we propose a novel cross-modal graph pre-training framework, called MIGA. By introducing phenotypic features of cellur images during pre-training, MIGA is able to bring generalisable biological priors to graph neural networks, enabling them to be transferred to downstream biologically relevant tasks. Experiments on a variety of downstream tasks demonstrate that our model outperforms SOTA machine learning-based methods and SSL methods, verifying the power of cross-modal pre-training strategy in the molecular graph representation area.

While our model shows promising results on graph-image representation learning, we have not yet dealt well with the issues of inherent data noise and batch effect. Thus, future directions include designing specific encoders for cellular images, optimizing cross-modal fusion mechanisms and introducing heterogeneous cross-modal data to improve the performance of graph pre-training.

## Acknowledgments and Disclosure of Funding

We would like to thank Minzhi Jiang and Wei Lu for discussion. This study has been supported by the National Key R&D Program of China (2020YFB020003), National Natural Science Foundation of China (61772566), Guangdong Key Field R&D Plan (2019B020228001 and 2018B010109006), Introducing Innovative and Entrepreneurial Teams (2016ZT06D211), Guangzhou S&T Research Plan (202007030010).

## A CIL-750K details

The original CIL dataset includes 919,265 five-channel fields of view containing 30,616 test compounds. It also includes metadata files which record morphological features for each cell in each image, both at the single cell level and at the population average level (i.e. per well); a workflow for image analysis to generate morphological features is also provided. Quality control indicators are provided as metadata, indicating fields of view that are out of focus or contain highly fluorescent material or debris. Chemical annotations are also provided for the application of compound processing. Figure 4 shows the molecular data distribution and the number of view per molecule in CIL dataset.

**Figure 4:**
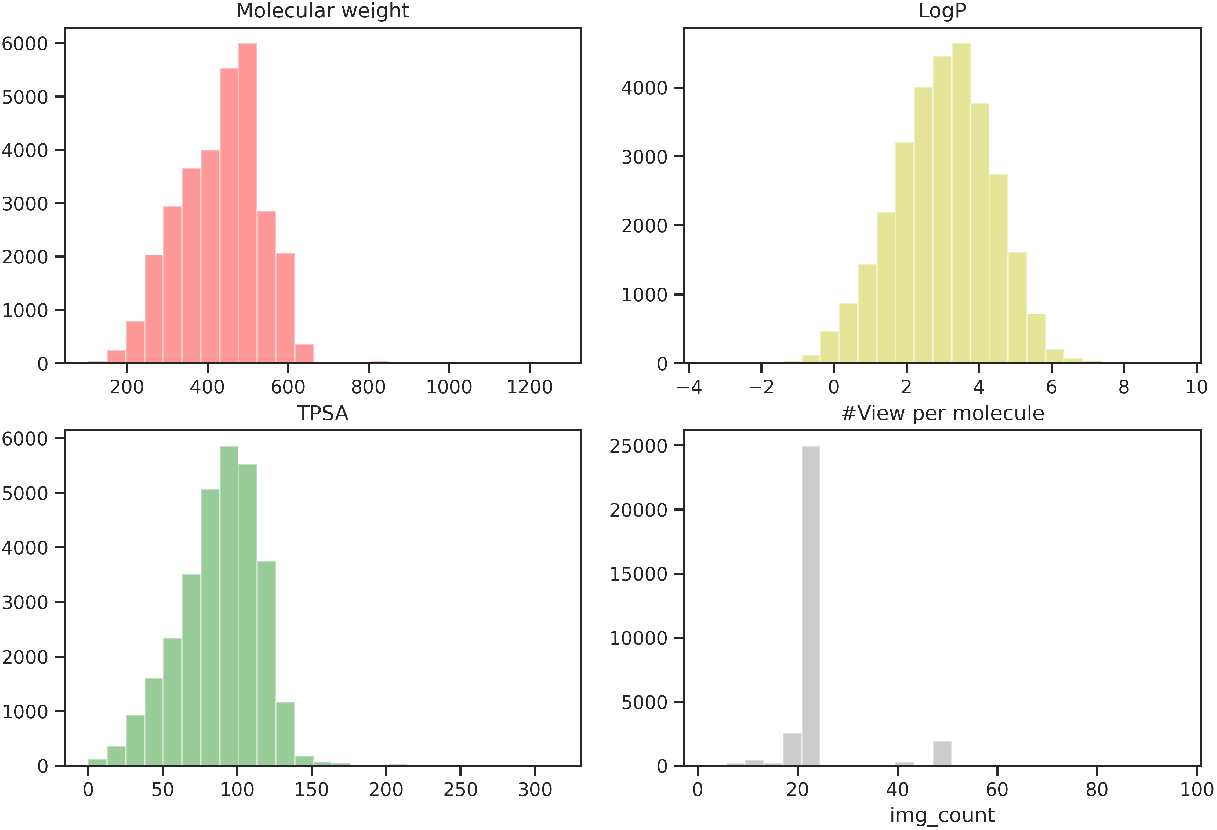
Data Distribution of CIL.

In CIL, each molecular intervention is imaged from multiple views in an experimental well and the experiment was repeated several times, resulting in an average of 30 views for each molecule. In order to keep the data balanced, we restricted each molecule to a maximum of 30 images and removed the untreated reference images, resulting in a cross-modal graph-image benchmark containing 750K views. Each view has a resolution of 692 × 520 pixels and 5 channels. These images were imaged with the ImageXpress Micro XLS automated microscope at 20× magnification. We resize the images to 128 × 128 without any cropping to fit the CNN models’ input format. Figure 5 shows examples of molecules and corresponding images from the CIL dataset and Figure 6 shows the multiple views of a random selected molecule.

**Figure 5:**
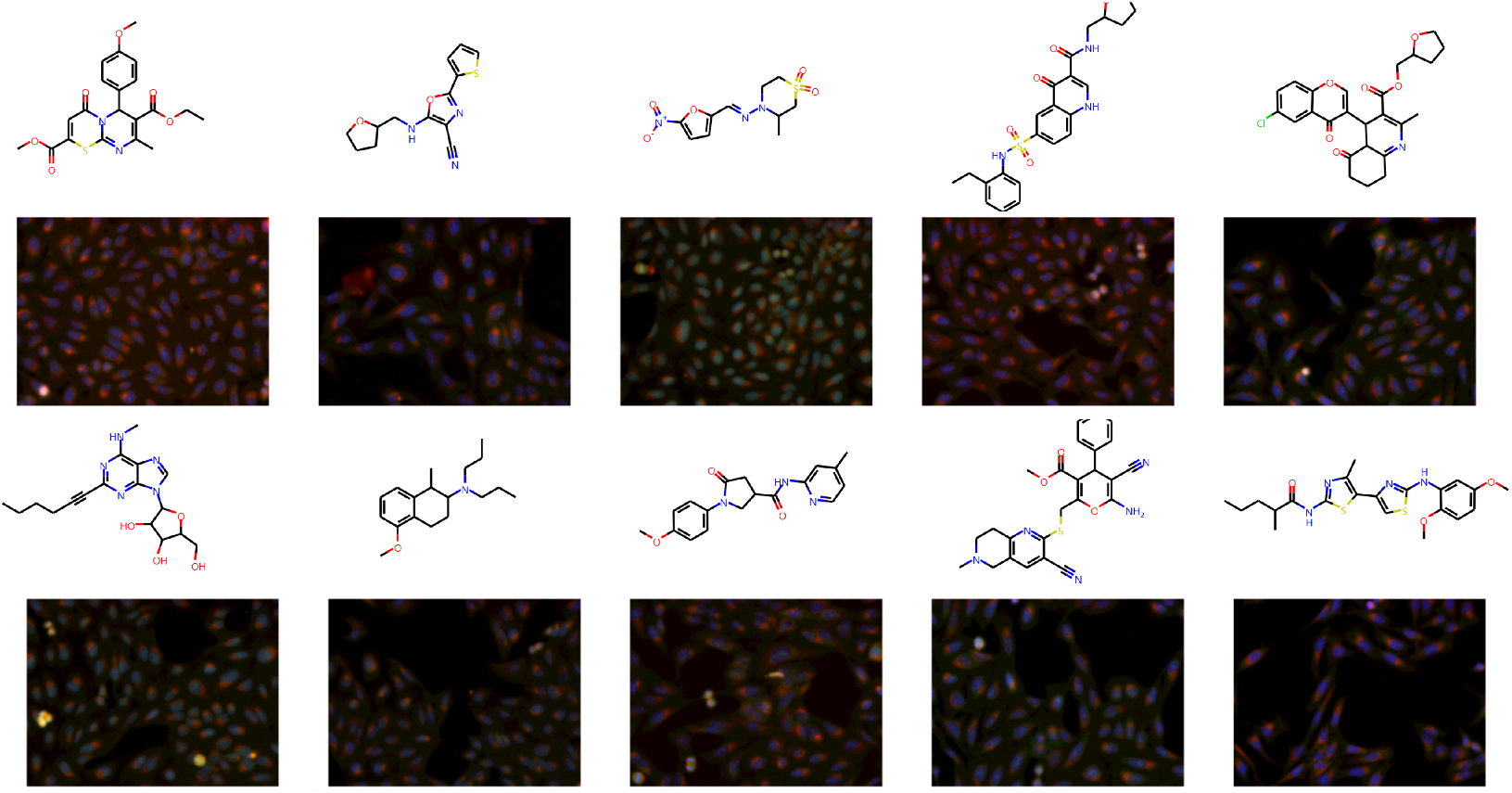
A random selection of 10 molecules and corresponding cellular images (1 view).

**Figure 6:**
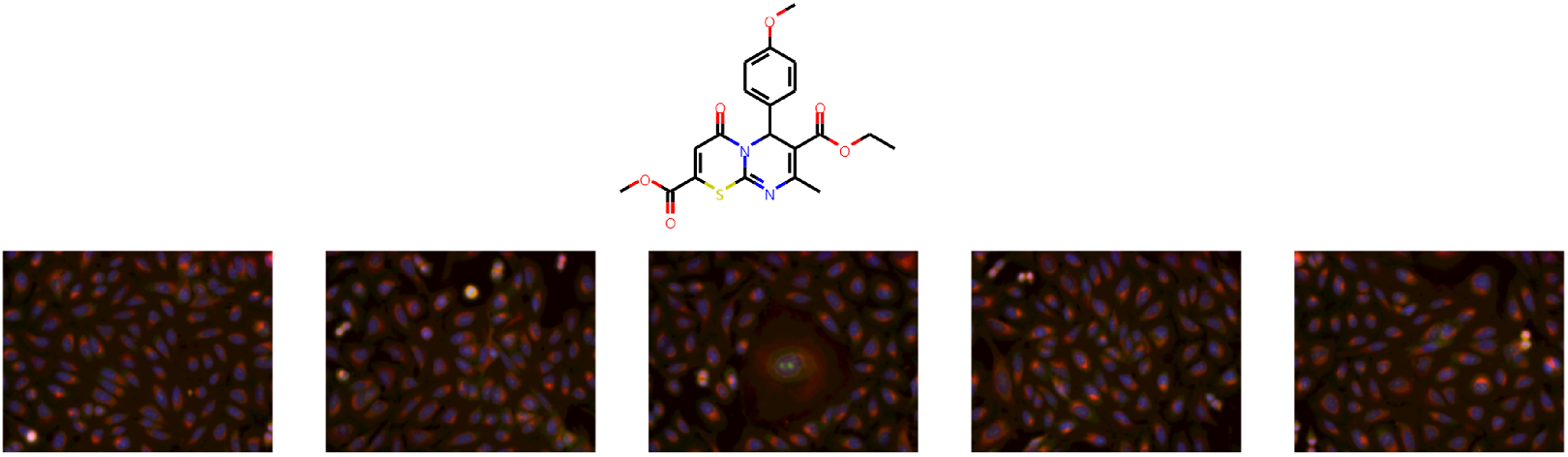
Five different views on the same molecule.

## B Implementation details and hyperparameters

Here we describe the implementation details for pre-training and fine-tuning stages.

### Pre-training

Our pre-training model consists of a Graph Isomorphism Network (GIN) from [3] with 5 layers and 300 hidden dimensions and a residual convolutional neural network (ResNet-34) [39] with 63.5M parameters. We pre-train the model for 100 epochs using a batch size of 1024 on 8 NVIDIA 3090TI GPUs. We use the Adam optimizer with an initial learning rate of 3e-4 and weight decay of 0.02. We take image with resolution of 128 × 128. The margin *γ* is set to 4. Pre-training on 750k graph-image pairs takes 4 hours, compared to 68 hours for GraphCL on 280k molecules of GEOM-Drugs and 110 hours for GROVER on

### Graph-Image Retrieval

We randomly split the CIL-750K dataset into a training set of 27.6K molecules corresponding to 680K images, and hold out the remaining of the data for testing. The held-out data consists of 3K molecules and the corresponding 50K images. We formulate the retrieval task as a ranking problem. In the inference phase, given a query molecular graph in the held-out set, we take images in the held-out set as a candidate pool and rank candidate images according to the L2 distance between the image embeddings and the molecular embeddings, and vice versa. The negative sampling rate is set to positive: negative = 1:100. We use GraphCL [12] and cross-modal pre-learning methods, CLIP [35] and ALIGN [36] as well as as baselines. The encoder part of these methods has been changed to the same setting as MIGA, but the decoder, training part and technical tricks have not been changed. After pre-training, We use the pre-trained model to output embeddings of molecular graphs and celluar images, then rank the candidate pool based on their L2 similarity.

### Zero-shot Graph Retrieval

We use MIGA to prioritize the functional molecules from a compound pool. This task mimics the real-world virtual screening scenario using morphological features observed when overexpressing a specific gene by cDNA interventions. We collected cellular images that were overexpressed with cDNA open reading frames for 6 genes by [53], including BRCA1, HIF1A, JUN, STAT3, TP53 and HSPA5. We used ExCAPEDB database [54] to retrieve gene-specific agonists and inactive molecules that have not been observed in training set. For each gene, we constructed a candidate pool with 20 agonists and 100 negative molecules, denoted as *P* (*I, G*_*A*_, *G*_*N*_). For each gene, given a random selected input image *I* with resolution of 128 × 128, we ask the model to rank the *G*_*A*_ in front of the *G*_*N*_. We use Hit@10 as the metric to evaluate our model in such a zero-shot graph retrieval task, where the random Hit@10 is 0.17 (20120).

### Clinical Outcome Prediction

#### Dataset

To standardize the clinical-trial-outcome predictions, we use the Trial Outcome Prediction (TOP) benchmark constructed by HINT, which incorporate rich data components including drug molecule information, disease information, trial eligibility criteria and trial outcome information. Herein, we consider phase-level evaluation on the trial outcome, where we predict the outcome of a single-phase study. Since each phase has different goals (e.g., phase I is for safety, whereas phases II and III are for efficacy), we evaluate phases I, II, and III separately. We follow the data splitting proposed by HINT and data statistics are shown in Table 6

**Table 6:** Statistics of Clinical Outcome Datasets.

| Phase-level | Molecule | Successes | Failures |
| --- | --- | --- | --- |
| Phase I | 944 | 564 | 380 |
| Phase II | 2865 | 1396 | 1469 |
| Phase III | 1752 | 1203 | 549 |

#### Baselines

We first include three machine learning-based methods (RF, LR, XGBoost) and a knowledge-aware GNN model HINT as our baseline. **Random Forest (RF)** is a bagging algorithm for classification or regression problems, which obtains the prediction by voting or averaging of each base learner (decision tree). **Logistic regression (LR)** is a simple, parallelizable classification method that uses maximum likelihood estimation for parameter estimation. **XGBoost**, also called an extreme gradient boosting tree, uses CART regression tree or linear classifier as a base learner to ensemble model predictions. These machine learning baselines utilize 1024-dimensional Morgan fingerprint features for trial outcome prediction. **HINT** is a hierarchical interaction network designed for clinical-trial-outcome predictions. It uses (1) 1024-dimensional Morgan fingerprint features, (2) a pre-trained BERT model to encode eligibility criteria into sentence embedding and (3) a graph-based attention model GRAM to encode disease information. Furthermore, we also include the self-supervised learning methods to constitute our baselines, including **ContextPred, GraphLoG, GROVER, GraphCL** and **JOAO**. For this downstream task, we use the molecule encoders over input molecule graphs for the fine-tuning of clinical outcome prediction.

#### Fune-tuning hyperparameter

For fine-tuning, we follow the GraphCL’s [12] settings. An extra linear classifier is appended to the pre-trained GNN. We fine-tune the model for 100 epochs using a batch size of 32 with a dropout rate of 50%. We use the Adam optimizer with an initial learning rate of 1e-3. Experiments are performed for 5 times, with mean and standard deviation of ROC-AUC and PR-AUC are reported.

### Molecular property Prediction Dataset BBBP

The Blood-brain barrier penetration dataset includes binary labels for 2035 compounds on their permeability properties.**Tox21**: The Tox21 dataset was created in the Tox21 data challenge, which contains qualitative toxicity measurements for 7821 compounds on 12 different targets, including nuclear receptors and stress response pathways. **HIV**: 41K compounds with binary labels for HIV virus replication inhibition. **ToxCast** includes 8576 drug compounds with binary labels of toxicity experiment outcomes with 617 targets. **ESOL**: The ESOL is a small dataset consisting of water solubility data for 1128 compounds.**Lipophilicity**: Experimental data for the octanol/water distribution coefficient of 4200 molecules.

#### Featurization Extraction

The feature extraction contains three parts: 1) Node feature extraction. 2) Bond feature extraction. 3) Topology connection matrix. We use RDKit to extract all features as the input of GNN. Table 8 and Table 9 show the atom and bond features we used in MIGA.

**Table 7:** Statistics of datasets. GC for Graph Classification, GR for Graph Regression.

| Dataset | Tasks | Type | Molecule | Metric |
| --- | --- | --- | --- | --- |
| BBBP | 1 | GC | 2,035 | ROC-AUC |
| Tox21 | 12 | GC | 7,821 | ROC-AUC |
| HIV | 1 | GC | 41K | ROC-AUC |
| ToxCast | 617 | GC | 8,576 | ROC-AUC |
| ESOL | 1 | GR | 1,128 | RMSE |
| Lipophilicity | 1 | GR | 4,198 | RMSE |

**Table 8:**
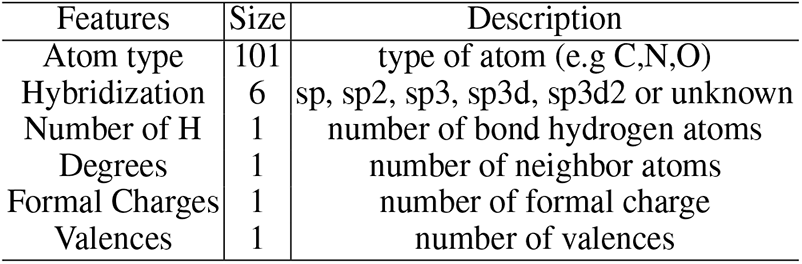
Atom features

**Table 9:** Bond features

| Features | Size | Description |
| --- | --- | --- |
| Bond type | 4 | single, double, triple, aromatic |
| Stereo | 6 | none, any, E/Z or cis/trans |
| In ring | 1 | whether the bond is part of a ring |
| conjugated | 1 | whether the bond is conjugated |

#### Fune-tuning hyperparameter

For fine-tuning, we followed the GraphCL’s [12] settings. An extra linear layer is appended to the pre-trained GNN to perform classification and regression, repectively. We fine-tune the model for 100 epochs using a batch size of 32 with a dropout rate of 50%. We use the Adam optimizer with an initial learning rate of 1e-3. Experiments are performed for 5 times, with mean and standard deviation of AUC and RMSE are reported.

## C Case study for image retrieval

**Figure 7:**
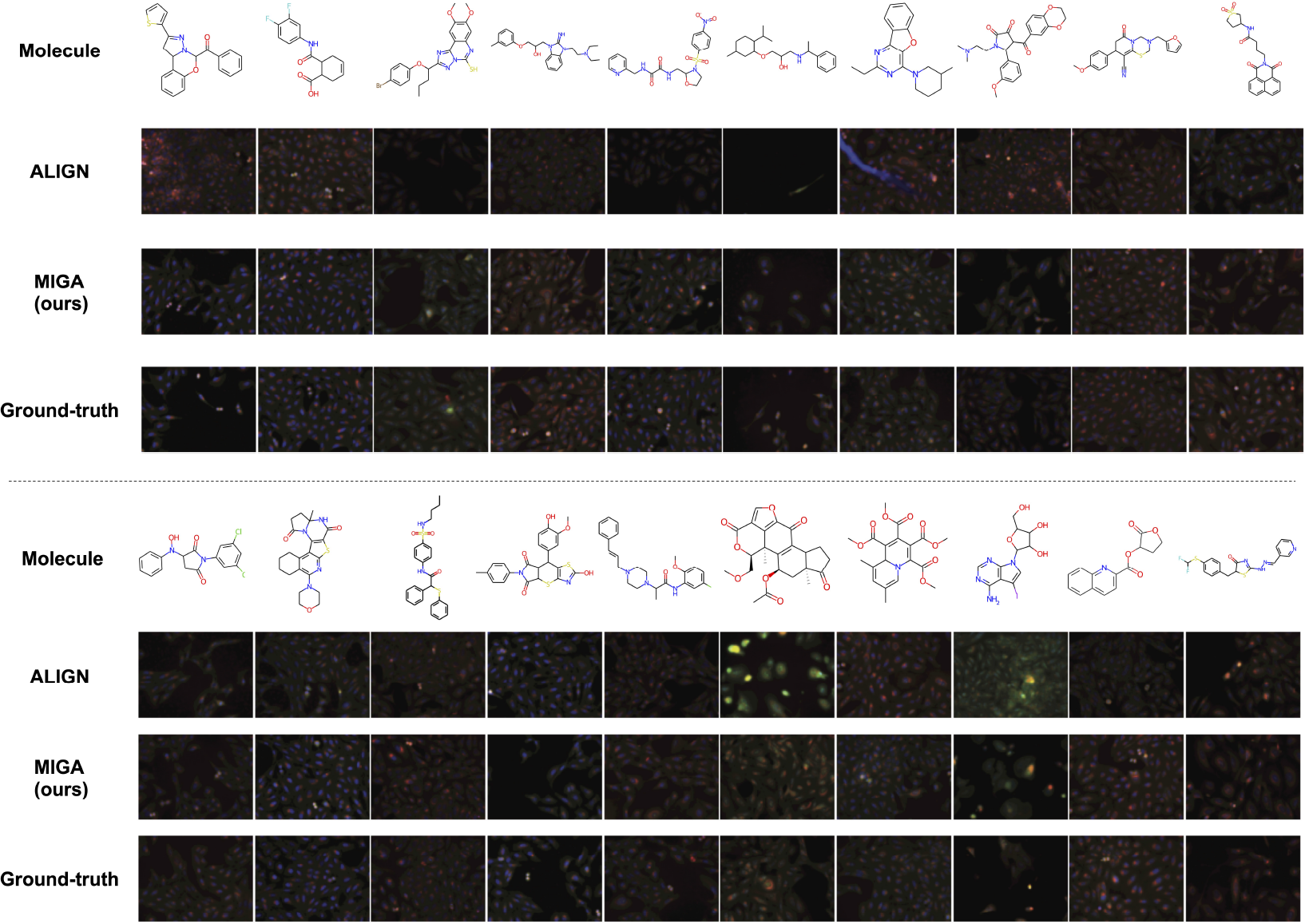
Case study for image retrieval task. The images retrieved by our method and baseline (ALIGN) are shown.

## D Case study for zero-shot graph retrieval

**Figure 8:**
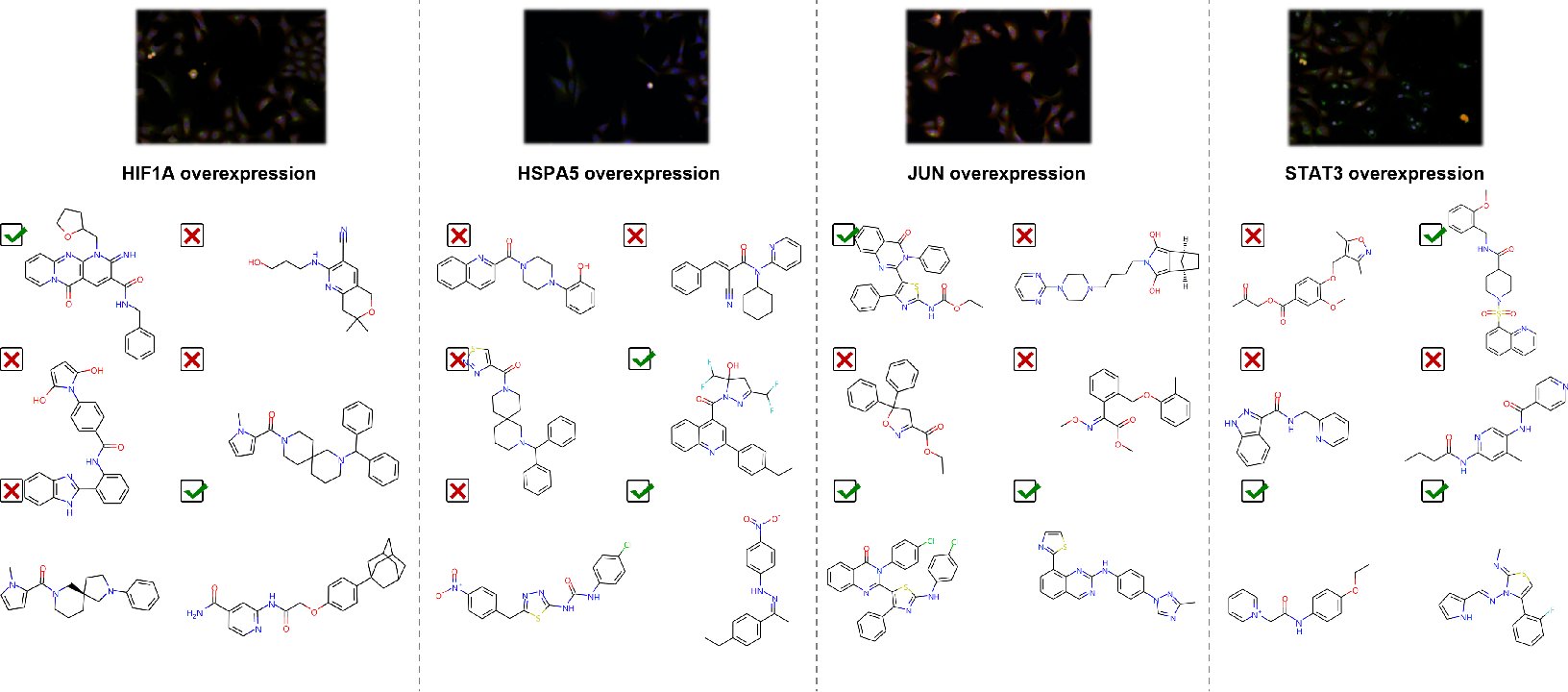
Case study for zero-shot graph retrieval task. The figure shows the cells induced by the cDNA interventions for specific genes (HIF1A, HSPA5, JUN, STAT3) and our model can find diverse molecules that have similar functions to these cDNA interventions (ticked).

